# Chrysalis: decoding tissue compartments in spatial transcriptomics with archetypal analysis

**DOI:** 10.1101/2023.08.17.553606

**Authors:** Demeter Túrós, Jelica Vasiljevic, Kerstin Hahn, Sven Rottenberg, Alberto Valdeolivas

## Abstract

Dissecting tissue compartments in spatial transcriptomics (ST) remains challenging due to limited spatial resolution and dependence on single-cell reference data. We present Chrysalis, a novel method to rapidly detect tissue compartments through spatially variable gene (SVG) detection and archetypal analysis without external references. We applied Chrysalis on ST datasets originating from various species, tissues and technologies and demonstrated state-of-the-art performance in identifying cellular niches.

## Main

ST is a rapidly evolving field dedicated to examining cells in their native tissue context, facilitating the identification of different cellular compartments or niches. Deciphering these elements is crucial, as they provide a substantial understanding of the tissue’s spatial structure, which largely determines its functionality^1^. A major obstacle in grid-based ST technologies, such as 10X Visium^2^ or Slide-seq^3^, lies in the precise identification of these tissue compartments and cellular neighbourhoods despite the limited spatial resolution^4^. To address this challenge, various computational methods have been developed for clustering^5–8^ and deconvolution^9–11^. Although these methods provide valuable insights, some limitations restrict their applicability and accuracy.

Unsupervised clustering methods have been adapted from single-cell RNA-sequencing (scRNA-seq) data analysis where each observation corresponds to a single cell^12^. Novel ST clustering methods have also incorporated spatial information^13^ and morphology^6^. However, the fundamental challenge remains categorising spots that simultaneously capture information from multiple cell types.

Alternatively, deconvolution methods aim to estimate the cell type composition by mapping reference signatures to the gene expression profiles obtained with ST. Although deconvolution methods present a robust alternative to clustering, they require representative scRNA-seq datasets, which are often not readily available^14^. Moreover, identifying new cell types presents an additional challenge for deconvolution methods, as they depend on the information defined in the reference dataset.

Techniques such as MEFISTO^15^ and NSF^16^ circumvent these constraints and offer insights into the tissue’s spatial architecture without requiring reference data by directly analysing factors from matrix decomposition models. However, the biological interpretability of their results and their computational efficiency is limited.

To address these limitations, we introduce Chrysalis, a novel computation method for the rapid detection of tissue compartments on grid-based ST datasets. Chrysalis employs archetypal analysis^17^, a method that has demonstrated its effectiveness in revealing the spatial organisation of breast cancer in a recent study^18^. Chrysalis identifies unique spatial compartments by archetypal decomposition of the low-dimensional representation derived from the SVG expression profiles (**Fig.1a**).

**Fig. 1.**
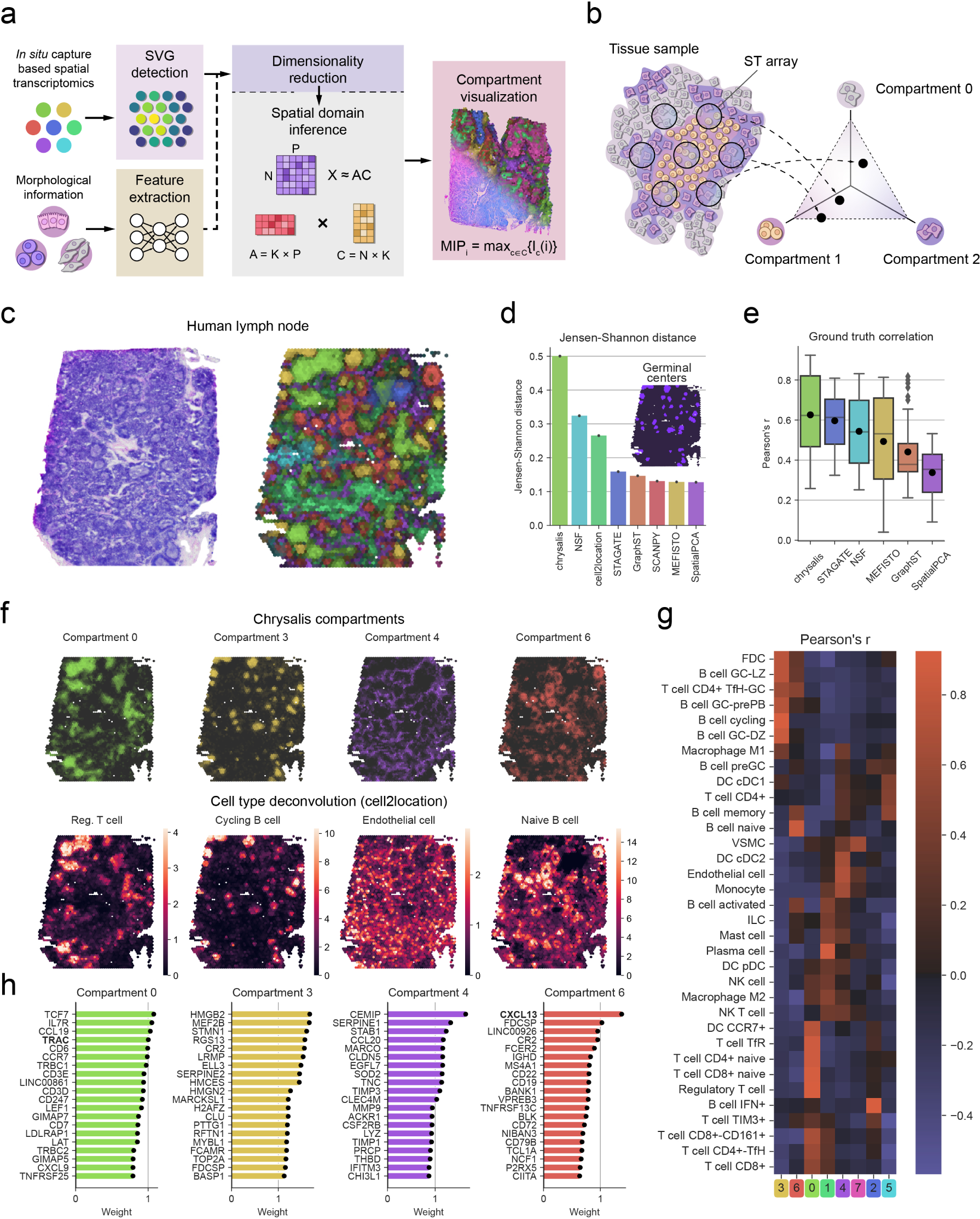
Chrysalis enables accurate tissue compartment identification in the human lymph node. **a**, Chrysalis workflow begins by selecting SVGs from the gene expression matrix to construct a low-dimensional representation of ST data using dimensionality reduction. This can be augmented by integrating morphological feature vectors extracted from the corresponding H&E image tiles. After dimensionality reduction, the integrated low-dimensional feature space is used to find spatial domains distinct in gene expression and, if available, morphology as well. By leveraging archetypal analysis, the original feature matrix *X* is decomposed into two matrices, *A* and *C*, where *C* contains the tissue compartment score for each observation, and *A* stores the contributions of the basis vectors for the tissue compartments. *A* can be used to reconstruct the weights of individual SVGs for *K* compartments. Finally, Chrysalis uses a maximum intensity projection-based visualisation to project *C* to the tissue space. **b,** Chrysalis finds discrete tissue compartments that appear as vertex points of a multi-dimensional simplex fitted to the low-dimensional feature space. These correspond to capture spots exclusively covered by one distinct cellular microenvironment, and every other capture spot containing the mixture of these compartments is represented as a linear combination of them. **c,** Human lymph node H&E image (left panel) and the projection of the tissue compartments identified by Chrysalis (right panel). **d,** Jensen-Shannon distance of domain score distributions in the germinal centre annotations (inset) and the rest of the sample across the applied methods. **e,** Boxplots of Pearson’s r values calculated between the 34 reference cell type abundances inferred with cell2location and the domain scores for the six methods applied to the lymph node dataset (centre line: median, box limits: upper and lower quartiles, whiskers: 1.5x interquartile range, diamonds: outliers, black dot: average value). **f,** Chrysalis Compartments 0, 3, 4, and 6 (upper panel) and their respective best-correlated cell type abundances (lower panel). **g,** Heatmap showing Pearson correlation coefficients between the cell type deconvolution results of the 34 reference cell types and the tissue compartments. **h,** 20 top contributing genes for Compartments 0, 3, 4, and 6. Bold characters were used to highlight the name of the genes referred to in the main text.

Chrysalis first identifies tissue compartments, i.e. archetypes, in the low-dimensional feature space that correspond to capture spots that are exclusively covered by a particular cellular niche (**Fig.1b**). Every other spot that contains signatures from multiple compartments is represented as a linear combination of these archetypes. In addition, the low-dimensional embedding of gene expression data can be easily combined with latent representations of morphological features extracted by deep learning models using the corresponding haematoxylin-eosin stained (H&E) image tiles. This integration enables the identification of compartments based on a unique combination of morphological characteristics and gene expression profiles. Furthermore, Chrysalis features a distinctive approach based on maximum intensity projection to visualise various tissue compartments simultaneously, facilitating the rapid characterisation of spatial relationships across the inferred domains. To demonstrate the capabilities of Chrysalis, we first examined a publicly available 10X Visium fresh frozen human lymph node sample. The lymph node exhibits intricate morphology, housing various populations of immune cells^19^, making it a challenging testing ground for spatial compartment detection. Chrysalis successfully distinguished tissue compartments that overlap with the primary morphological features in the histological image highlighted by the pathologist annotations (**Fig.1c and Supplementary Fig.1a**). Further using annotations, we benchmarked Chrysalis against other recently proposed methods aiming to identify spatial domains using various computational models^9,15,16,20–23^. Here, Chrysalis achieved the highest differential in spatial domain score distributions inside and outside the annotated germinal centres, showing its enhanced ability to distinguish microenvironments (**Fig.1d and Supplementary Fig.1b-e**).

Subsequently, we evaluated the biological relevance of the tissue compartments identified by Chrysalis (**Supplementary Fig.2**) by investigating their correlation with the abundance of 34 reference cell types mapped onto the lymph node sample using the cell2location^9^ cell type deconvolution method (**Fig.1e-g and Supplementary Fig.2a**). We assessed the performance of competing methods by calculating the average Pearson correlation coefficient between the deconvolution results and their corresponding outcomes. Chrysalis outperformed existing approaches, most notably the two main matrix decomposition-based techniques, NSF and MEFISTO (**Fig.1e**). Specifically, B cell populations predominating in germinal centres, such as cycling, GC-DZ, and GC-LZ cells, are associated with Compartment 3 with 0.91, 0.84 and 0.77 Pearson’s r, respectively. Naive B cells residing in B cell follicles matched with Compartment 6 (0.84 Pearson’s r). The overall tissue vasculature correlated with Compartment 4 (0.72 Pearson’s r with endothelial cells), and various T cell populations (naive, CD4+/CD8+, regulatory, TfR) showed a strong association with Compartment 0 with an average Pearson’s r of 0.89 (**Fig.1g**). As an orthogonal validation, we evaluated the correlation between the compartments and the expression of marker genes of lymph node resident immune cells (**Supplementary Fig.2c**). We found strong associations, confirming that these signatures can aid in interpreting the detected tissue compartments. Moreover, by directly examining the weights assigned to individual genes for each tissue compartment, we identified numerous canonical markers of different immune cells, such as *TRAC* or *CXCL13* (**Fig.1h and Supplementary Fig.2e-f**). This approach holds significant potential for characterising tissue compartments when prior knowledge is lacking.

Next, we applied Chrysalis to a human breast cancer dataset containing serial sections of a formalin-fixed paraffin-embedded (FFPE) ER+ / HER2+ tumour sample, analysed with 10X Visium and 10X Xenium^24^. Running Chrysalis on the Visium data revealed distinct tissue compartments exhibiting unique morphological and transcriptional features (**Fig.2a and Supplementary Fig.3**). Notably, these regions coincide with the pathologist’s annotations (**Supplementary Fig.4a**) of the invasive tumour region and multiple phenotypes of ductal carcinoma in situ (DCIS). The high resolution of Xenium is suitable for subcellular transcript detection, and combined with imaging-based cellular segmentation, enables single-cell phenotyping. We co-registered and transformed the Xenium readouts with the Visium spots to generate a reference for validating the tissue compartments inferred by Chrysalis (**Supplementary Fig.4b-d**). Using this reference as ground truth, we again conducted the correlation-based benchmark to evaluate the performance of different methods on this dataset, with Chrysalis achieving the highest mean Pearson’s r value (**Fig.2b**). The correlation between the compartments and the reference cell type-specific abundance data also indicated that the immune microenvironment (Compartment 5), the invasive/proliferative tumour (Compartment 2), the two separate DCIS phenotypes (Compartment 0 and 6), and the myoepithelial tissue (Compartment 4) were precisely identified by Chrysalis (**Fig.2c-d**). Compartments 1, 3, and 7 did not show significant correlations with any of the cell types defined by the DNA probe set, which lacks the capacity to differentiate distinct stromal cell types or adipocytes. However, by examining these compartments (**Supplementary Fig.3b**), we observed that Compartment 1 contained genetic markers of cancer-associated fibroblasts, such as *MMP2*, *COL1A1*, and *HTRA1*^25^. Compartment 3 included *FABP4*, *GPX3*, and *PLIN1*, which are highly expressed in adipocytes, and the top genes in Compartment 7 - *IGHG1*, *SFRP4*, and *PLTP* - are associated with tumour-infiltrating plasma cells^26,27^. In addition, inspecting capture spots with high domain scores also enabled us to calculate the cell type composition of each tissue compartment, demonstrating various cellular niches dominated by distinct tumour cell populations (**Supplementary Fig.3e**). The results of Chrysalis were further validated by assessing the weights of canonical tumour and myoepithelial markers (**Supplementary Fig.3f**) and the top compartment-specific genes (**Fig.2e**).

**Fig. 2.**
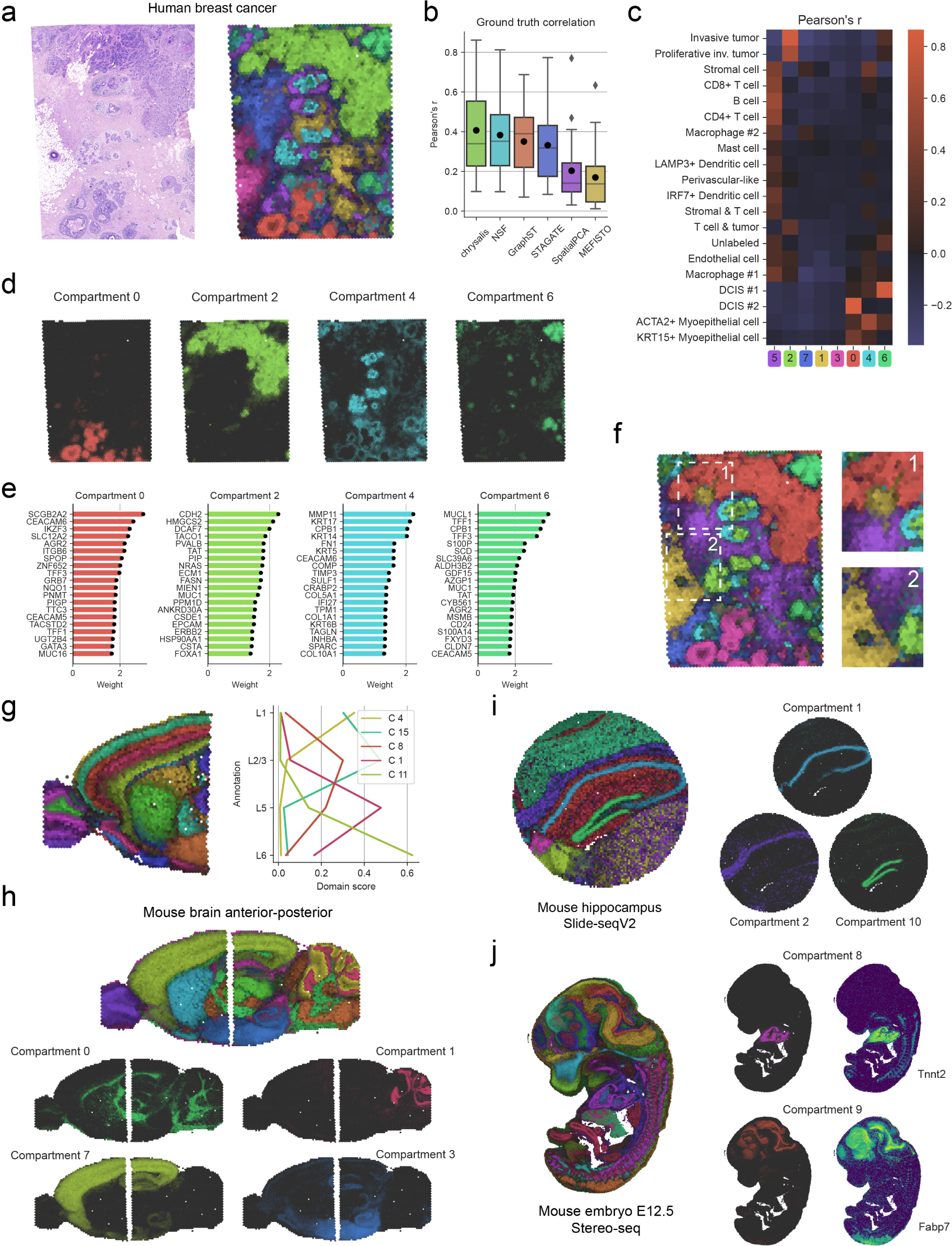
Chrysalis uncovers distinct tissue compartments across various tissue types and spatial technologies. **a,** Human breast cancer H&E image (left panel) and the projection of the tissue compartments identified by Chrysalis (right panel). **b,** Boxplots of Pearson’s r values calculated between the reference cell type abundances from the Xenium measurement and the domain scores for the six methods applied to the human breast cancer dataset (centre line: median, box limits: upper and lower quartiles, whiskers: 1.5x interquartile range, diamonds: outliers, black dot: average value). **c,** Correlation heatmap showing Peason’s r between the ground truth cell type abundance data and the identified compartments. **d,** Individual Chrysalis Compartments 0, 2, 4, and 6 denoting DCIS #1, invasive tumour, myoepithelial, and DCIS #2 signatures, respectively. **e,** Top 20 genes that contribute the most to Compartments 0, 2, 4, and 6 in the human breast cancer dataset. **f,** Chrysalis compartments with integrated morphological features extracted with a deep autoencoder (left panel). Morphological features improved the separation between the adipose tissue (yellow), invasive tumour (red), and stroma (purple; right panels). **g,** Tissue compartments within the mouse brain identified by Chrysalis (left panel). Mean domain scores in expert-annotated cortical regions (right panel). **h,** Chrysalis composite plot of the two-sample mouse brain dataset (upper panel) and individual compartments corresponding to (left-to-right then top-to-bottom) the fibre tracts, internal granular layer of the cerebellum, and ventral and dorsal cortex. **i,** Chrysalis projection of the mouse hippocampus captured with Slide-seqV2 (left panel) and individual compartments showcasing distinct cellular microenvironments (right panel): pyramidal layer of the Ammon’s horn (Compartment 1), corpus callosum (Compartment 2), and molecular layer of the dentate gyrus (Compartment 10). **j,** Projection of Chrysalis compartments in the mouse embryo (E12.5) captured with Stereo-seq (left panel) and the compartments corresponding to the developing heart and brain alongside their corresponding top-weighted genes.

As histomorphology is usually well conserved in FFPE material, we further utilised this sample to illustrate the potential of incorporating morphological features into Chrysalis. To achieve this, we trained a deep autoencoder on the H&E image tiles corresponding to each capture spot and integrated the encoder output with the gene expression data, creating a multimodal representation. With this approach, the inferred tissue compartments align more closely to the overall tissue morphology and our pathologist’s annotations (**Fig.2f and Supplementary Fig.5**). Furthermore, we measured how well the morphology-integrated tissue compartments overlap with the annotations compared to the gene expression-based compartments. We observed an increase in Adjusted Rand Index (ARI) and F_1_ scores, e.g.: F_1_ score for adipose tissue increased from 0.60 to 0.87 (**Supplementary Fig.5b-c**).

Subsequently, we analysed a 10X Visium dataset of a fresh frozen parasagittal mouse brain tissue section with Chrysalis. The brain tissue exhibits a complex arrangement of layers in the neocortex, making it challenging to accurately recapitulate the exact anatomical structures. The tissue compartments were assessed by a trained veterinary pathologist, using the olfactory bulb and the Allen Mouse Brain Atlas^28^ as references. Chrysalis precisely identified the 6 layers of the olfactory bulb, the accessory olfactory bulb, the layers of the somatomotor cortex, striatal structures, the hippocampus CA3 region as well the thalamus, pallidum and hypothalamus (**Fig.2g left panel and Supplementary Fig.6a**). These results were supported by the increased domain scores within the cortical layer annotations (**Fig.2g right panel and Supplementary Fig.6b**). Genes unique to each compartment revealed cell-type markers and key contributors to each cortical layer, supported by the mouse brain transcriptional atlas^29^ (**Supplementary Fig.7**). For instance, genes such as *Ptgds* and *Mgp*, markers for leptomeningeal cells and excitatory neurons respectively, were found in Compartment 4 (layer 1). Compartment 15 is marked by *Calb1* and *Lamp5*, which are enriched in layer 2/3. Meanwhile, *Vxn*, a marker for layers 6a-b, can be found in Compartments 1 and 11.

In addition, we used this sample to showcase the performance of Chrysalis on datasets containing multiple tissue samples by combining it with the posterior section of the mouse brain. On this integrated tissue sample, Chrysalis readily identified the main anatomical regions of the brain, including the Ammon’s horn and the dentate gyrus (**Fig.2h top panel and Supplementary Fig.8**). By further investigating the individual tissue compartments (**Fig.2h bottom panels and Supplementary Fig.8b**), we observed continuous compartments that extend across both tissue sections. As an example, Compartment 0 corresponds to the fibre tracts with top genes such as *Plp1, Mbp,* and *Mobp* marking oligodendrocytes. In contrast, Compartment 1 can be exclusively found in the posterior brain sample, highlighting the internal granular layer of the cerebellum as indicated by the cerebellin genes. Compartments 7 and 3 are in accordance with the dorsal and ventral cerebral cortex, respectively (**Supplementary Fig.8d**).

We demonstrated the versatility of Chrysalis by applying it to datasets acquired with Stereo-seq^30^ and Slide-seqV2 ST platforms. In a mouse hippocampus dataset generated with Slide-seqV2, Chrysalis distinguished the pyramidal layer of the Ammon’s horn (Compartment 1) and delineated the molecular layer of the dentate gyrus (Compartment 10) from the surrounding brain tissue (**Fig.2i and Supplementary Fig.9**). Moreover, Chrysalis pinpointed tissue compartments in mouse embryo cross-sections (E9.5 and E12.5) measured with Stereo-seq, corresponding to the developing organs, such as the heart (Compartment 8), liver (Compartment 21) or brain (Compartment 9), corroborating the expert annotations from the original study (**Fig.2j and Supplementary Fig.10**).

In summary, we present Chrysalis, an innovative method that employs SVG identification and archetypal analysis for rapid tissue compartment identification and visualisation in ST with state-of-the-art performance. Additionally, Chrysalis is applicable to any grid-based ST technology and can be expanded by integrating morphological features. We anticipate that Chrysalis will emerge as a valuable extension of ST data analysis workflows, enhancing our understanding of the spatial structure and functional characteristics of tissues, thus propelling the field of ST further into the realm of high-resolution biological discovery.

## Methods

### Chrysalis

#### Data preprocessing

We have designed Chrysalis to seamlessly integrate into workflows based on SCANPY^23^ or Squidpy^31^ Python packages. As such, SCANPY functions are utilised for quality control (QC) and preprocessing for all datasets. Given that Chrysalis assumes that unique tissue compartments are represented by extreme points, stringent QC and outlier removal are important for finding high-quality tissue compartments. During our analyses, SCANPY functions were used to perform QC by first filtering spots based on the unique molecular identifier (UMI) count numbers (*pp.filter_cells*). The exact cutoff value can be determined individually for samples based on the UMI count distribution and other technical attributes. Genes expressed in less than 10 capture spots (*pp.filter_genes*) were subsequently discarded. Following that, the gene expression matrix was normalised (*pp.normalize_total*) and log1p-transformed (*pp.log1p*) using SCANPY preprocessing functions before subjecting it to further analysis.

#### Detection of SVGs

PySAL’s^32^ implementation of the global Moran’s Index is used (**Supplementary Fig.11**) to detect SVGs in the gene expression matrix. First, we standardise the preprocessed (normalised and log1p-transformed) expression matrix so that for each capture spot *i*, the standardised gene expression *z*_*i*_ can be described as:

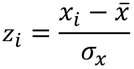

where *x*_*i*_ represents the expression level of a specific gene at the *i*-th capture spot, while 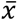 is the mean expression level across all spots and σ_*x*_ is the standard deviation of expression levels across all spots. Moran’s I^33^ can then be formulated as follows:

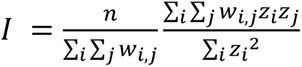

where *z*_*i*_ and *z*_*j*_ are the standardised gene expression levels at spots *i* and *j*, respectively. *n* is the total number of spots and *w*_*ij*_ is the spatial weight between spot *i* and spot *j*. Spatial weight *w*_*ij*_ is defined as 1 if spot *j* is within the set number of k-nearest neighbours (KNN) of spot *i* and 0 otherwise. To evaluate the potential influence of neighbourhood size, we computed Moran’s I statistic with 6, 18, and 36 neighbours (**Supplementary Fig.11c**) on the 10X Visium human lymph node sample (for details on preprocessing, see **Methods, Human lymph node analysis** section). As our results show consistent Moran’s I values regardless of neighbourhood size, Chrysalis uses 6 nearest neighbours by default for computational efficiency, reflecting the hexagonal grid pattern in which Visium capture spots are organised. For analysing Stereo-seq data, we used a neighbourhood size of 8, as the DNA nanoball particles are arranged according to the Cartesian coordinate system. The Slide-seqV2 dataset was analysed with the same neighbourhood size of 8 due to the preprocessing we performed (see **Methods, Slide-seqV2 analysis** section).

Moreover, we also explored alternative spatial autocorrelation statistics instead of Moran’s I, namely, Geary’s C^34^. It has been previously reported^35^ that combining the two metrics can yield better performance, as Geary’s C is more sensitive to local changes. Nevertheless, these two metrics were indistinguishable in our tests using the preprocessed 10X Visium human lymph node dataset (for details on preprocessing, see **Methods, Human lymph node analysis** section), and therefore we continued with Moran’s I (**Supplementary Fig.12a**). To decrease computational runtimes, Chrysalis calculates Moran’s I only for genes that are expressed in at least 5% of the capture spots. This approach does not compromise the identification of spatially variable genes, as demonstrated by the 2D histogram showing that genes exhibiting high Moran’s I values tend to be expressed across wider regions of the tissue (**Supplementary Fig.11b**).

To determine the number of SVGs, a rank-order plot is generated by arranging the Moran’s I values for every gene in descending order. This results in a curve characteristic of a power-law distribution (**Supplementary Fig.11d**). The inflection point of this curve, marking the transition from a high-autocorrelation interval to a flatter low-autocorrelation interval, serves as a cutoff point for classifying SVGs. Genes below this inflection point are omitted, preserving most of the spatial variability in the data while reducing noise. Our findings indicate that in the majority of cases, the inflection point occurs approximately at the 1000th gene. Consequently, Chrysalis classifies the top 1000 genes as SVGs by default, which are then used for further analysis.

For SVG detection, Chrysalis can also incorporate alternative methods^36–39^, which may provide marginal improvements in performance at the cost of increased computational runtimes (**Supplementary Fig.12b-d**). To compare the performance of these SVG detection methods to Moran’s I, the 10X Visium human lymph node dataset was first obtained from the 10X Genomics online data repository and preprocessed using SCANPY. SVG detection methods SpatialDE2^36^, SEPAL^37^, BSP^38^, and SPARK (SPARK-X)^39^ were run according to the published guidelines (**Supplementary Table S1**). Chrysalis was then applied using the top 1000 SVGs assigned by each method (for details on preprocessing, Chrysalis pipeline, and correlation-based validation, see **Methods, Human lymph node analysis** section). Pearson correlation coefficient was then calculated between the tissue compartments and the cell type deconvolution results.

#### Low-dimensional embedding

Principal component analysis (PCA) is performed (Scikit-learn^40^ Python package implementation) on the SVG matrix to identify the main spatial expression patterns. To determine the optimal number of PCs, we conducted multiple tests with varying configurations using the human lymph node dataset (PCs: [6, 10, 20, 40], compartments: 3-24). We calculated a correlation matrix between the cell type deconvolution results and the compartments identified by Chrysalis (see the details in **Methods, Human lymph node analysis** section) and assessed the average Pearson’s r value of the highest correlating cell type-tissue compartment pairs to evaluate the compartments’ ability to depict the true tissue microenvironments (**Supplementary Fig.2g**). Based on this analysis we set the default to 20 PCs, as it sufficiently captured information attributed to the underlying tissue microenvironments.

#### Morphological feature integration

Chrysalis can incorporate latent feature vectors obtained through deep learning models from H&E images associated with ST measurements. This allows the identification of tissue compartments with unique transcriptional and morphological properties. The generic workflow includes:

1. Extracting morphological features for each capture spot
2. Performing dimensionality reduction on the feature vectors
3. Concatenating the morphology-based and gene expression-based components
4. Scaling the concatenated components to prevent any issues arising from disparate scales across modalities

The integrated feature matrix can be subsequently used in Chrysalis to find tissue compartments with archetypal analysis (for details regarding the integration performed for the human breast cancer data, see **Methods, Human breast cancer analysis, Morphological feature extraction and integration** section).

#### Tissue compartment inference

To detect tissue compartments, Chrysalis uses archetypal analysis on the low-dimensional embedding of SVGs. The aim of archetypal analysis is to find extremal points in the multidimensional data known as archetypes (i.e. tissue compartments) such that any sample from the dataset can be represented as a convex combination of these archetypes. This can be interpreted as fitting a multidimensional simplex to the data space with the archetypes representing the vertices of this simplex (**Fig.1b and Supplementary Fig.2d**). Therefore, any observation in the data (e.g. each capture spot) can be represented as a mixture of archetypes with a measurable contribution of each archetype to the sample which allows for a more comprehensive understanding of the tissue structure.

More formally, let *X* be a data matrix of size *N* × *P*, where *N* is the number of observations (capture spots) and *P* is the dimension of low-dimensional SVG representation (selected number of principal components).

Archetypal analysis aims to find a set of *K* archetypes such that the input data matrix *X* ∈ *R*^*N*×*P*^ can be approximated as *X* ≈ *CA*, where *C* ∈ *R*^*N*×*K*^ denotes the coefficient matrix and *A* ∈ *R*^*K*×^ ^*P*^ denotes archetypes. This approximation is subject to the following constraints:

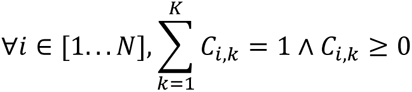

Where *C*_*i*,*k*_ denotes the contribution of *k*-th archetype to the *i*-th observation.

The process of obtaining matrices *C* and *A* involves minimising the reconstruction error, measured as the Frobenius norm across all data points formulated as follows:

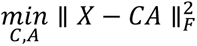

Moreover, each observation is approximated as:

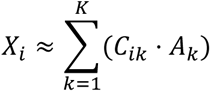

Upon optimisation, the low-dimensional SVG representation is decomposed into *K* archetypes. In order to obtain archetypical decompositions of SVGs, the PCA loadings can be used as described below. Let *L* represent the loadings obtained through PCA decomposition of SVGs. The expression of the individual genes for archetype *k* represented by *E*_*k*_ can be calculated as follows:

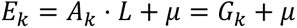

where μ is the preprocessed mean (normalised and log1p-transformed, for details, see **Methods, Data preprocessing** section) gene expression vector, and *G*_*k*_ represents the weights of each gene in the *k*-th archetype.

Chrysalis uses the *archetypes* Python package implementation of archetypal analysis (n_init=3, max_iter=200, tol=0.001, random_state=42).

#### Visualisation

To jointly visualise the tissue compartments, maximum intensity projection (MIP) is used. Initially, each archetype *k* (tissue compartment) is assigned to a unique colour *I*_*k*_ in the RGB space, with each colour channel containing values in the range [0, 255]. Following that, linear interpolation is used to assign colours to each capture spot *i* for every tissue compartment *k*. Consequently, the assigned colour values transition between (0,0,0), which is assigned to the lowest value in compartment *k*, and *I*_*k*_. Given *K* tissue compartments, the visualisation colour for a spot *i* is determined using MIP as follows:

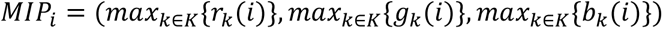

where *r*_*k*_(*i*), *b*_*k*_ (*i*), and *b*_*k*_ (*i*) represent the red, green, and blue values for compartment *k* at spot *i*.

### Human lymph node analysis

We obtained the 10X Visium human lymph node dataset from the 10X Genomics online data repository and preprocessed it using the SCANPY package as described below. First, capture spots containing less than 6000 UMI counts were removed (*pp.filter_cells*) to filter out low-quality spots. Genes expressed in less than 10 spots were discarded (*pp.filter_genes*). Next, we normalised (*pp.normalize_tota*l) and log1p-transformed (*pp.log1p*) the gene expression matrix. We then employed Chrysalis to identify SVGs using Moran’s I, selecting genes with at least 0.08 Moran’s I value amongst genes expressed in more than 5% of the capture spots. The included histology image was used to annotate tissue types. 997 SVGs above the previously defined Moran’s I threshold were used to construct a low-dimensional representation using the dimensionality reduction step of Chrysalis. Finally, Chrysalis was used to define tissue compartments using archetypal analysis (number of PCs=20, number of compartments=8).

### Validation with cell type deconvolution results and specific marker genes

#### Correlation with reference data

Cell type deconvolution results were generated with cell2location^9^ by running the Google Colaboratory notebook provided in the documentation (**Supplementary Table S1**), using scRNA-seq reference data from the original publication. This single-cell reference was originally obtained by integrating datasets from human secondary lymphoid organs comprising more than 70,000 cells. Correlation coefficients were calculated using the Pearson method to analyse the relationships between the 8 tissue compartments identified with Chrysalis and the inferred abundances of 34 cell types. Subsequently, these coefficients were used to construct a correlation matrix for validating the tissue compartments.

#### Correlation with marker gene signatures

We explored canonical marker gene signatures of 16 cell types found in the lymph node derived from the CZ CELLxGENE Discover online resource. Using SCANPY’s *tl.score_genes* function, scores for each of the 16 downloaded gene sets were determined by calculating the difference between their average expression and the average expression of a reference gene set, which was randomly sampled from the total gene pool corresponding to discretised expression levels. Pearson correlation coefficients were calculated, and a correlation matrix was constructed to denote the association between the signature scores and the compartments.

### Benchmarking

#### Calculating low-dimensional embedding of alternative methods

We first preprocessed and analysed the lymph node dataset with several recently published spatial domain detection methods offering alternative low-dimensional embeddings or corrections of the gene expression matrix by various machine learning and deep learning models. For NSF^16^ and MEFISTO^15^ the gene expression matrix was decomposed to 8 factors using their respective software implementation pipelines (**Supplementary Table S1**) after preprocessing the data with SCANPY following the same criteria as for Chrysalis. For STAGATE^20^, GraphST^21^, and SpatialPCA^22^, the low-dimensional embeddings were extracted after running their respective inference pipelines according to the published guidelines (see **Supplementary Table S1**). This resulted in low-dimensional embeddings with 30 components for STAGATE and 20 for SpatialPCA. As GraphST outputs the corrected gene expression matrix, we applied a PCA and used the first 6 PCs for benchmarking. These represent 74% of the total explained variance, corresponding to the point of inflection in the cumulative explained variance curve.

#### Assessment of alternative methods in characterising germinal centres

To benchmark the effectiveness of the low-dimensional representations in characterising the annotated germinal centres, we selected spatial domains (i.e. individual factors from NSF and MEFISTO, embedding vectors from STAGATE, and SpatialPCA, and PCs for GraphST) best describing the annotated germinal centres as described below. First, the low-dimensional representations were min-max normalised to be within the range of 0 to 1. Subsequently, using the germinal centre annotations from the cell2location article^9^, we calculated ROC-AUC scores to identify the embedding vectors most representative of the germinal centres. (**Supplementary Fig.1b-c**). In addition, we added the combined abundance of specific cell types found in the germinal centres (B cell GC-DZ, B cell GC-LZ, B cell cycling, T cell CD4+ TfH-GC, B cell GC-prePB, FDC) that were deconvolved using cell2location to the comparison along with a PC calculated with the default SCANPY pipeline, based on highly variable gene detection (*pp.highly_variable_genes*) and SCANPY’s PCA function (*pp.pca*). Following that, the mean Jensen-Shannon distance was calculated between the embedding vector (i.e. domain score) distributions (**Supplementary Fig.1d**) for spots labelled as germinal centres and the rest of the sample from 1000 random permutations.

#### Benchmarking alternative methods with cell type deconvolution results

For the correlation-based benchmark, the unscaled low-dimensional representations were used. Correlation matrices were first built by calculating Pearson correlation coefficients between the 34 cell type abundances from the cell2location reference data and the embeddings. For each cell type, the highest Pearson’s r value was selected, corresponding to the highest correlating embedding vector, and the averages of these values were calculated.

#### Computational running time

The computational running time of these methods was gauged by measuring execution time starting by reading the dataset up to the point where the low-dimensional representations were calculated (**Supplementary Fig.1e**). This measurement was done on a Windows machine using Ubuntu 20.04 LTS via Windows Subsystem for Linux without GPU acceleration (CPU: AMD Ryzen 9 5900HS, RAM: 32 GB, GPU: NVIDIA GeForce RTX 3060 Laptop GPU).

### Human breast cancer analysis

The combined 10X Xenium and Visium human breast cancer dataset was downloaded from the 10X Genomics online resource with the corresponding high-resolution H&E. The Xenium dataset contains 167,780 segmented cells with the gene expression data of 313 genes. Cell type annotations of 20 cell types were directly provided by 10X Genomics (**Supplementary Fig.4b**).

#### Co-registration of Visium and Xenium data

To map individual cells from the Xenium section to the Visium data, we first co-registered the two corresponding H&E tissue images. We utilised the image alignment functionality of 10X Genomics’ Loupe Browser, which offers manual landmark-based image alignment. We placed 30 landmarks to create the initial alignment, followed by an algorithmic refinement step relying on the mutual information measured between the two images. Two image registration steps were carried out: one between the post-Xenium H&E and the image captured by the CytAssist machine, and another one between the pre-Visium H&E and the CytAssist image, as the image alignment functionality in Loupe Browser is limited to CytAssist generated image data. Afterwards, affine transformation matrices were exported and used to translate the XY positions of individual cells in the Xenium dataset to the Visium coordinate system (**Supplementary Fig.4c**). Cells with centroids outside of the Visium capture spots were omitted and the remaining ones were aggregated for each cell type and assigned to the capture spots (**Supplementary Fig.4d**). As there is no one-to-one mapping between the two modalities, spots outside of the Xenium ROI were also removed, retaining 3906 of the original 4992 capture spots.

#### Mapping pathologist annotations to the Visium dataset

The high-resolution post-Xenium H&E image was assessed and annotated by a trained veterinary pathologist using QuPath^41^ software. Transformation matrices described in the above paragraph were used to translate the annotation polygons and the spatial coordinate pairs of the Visium dataset to a unified coordinate system. Visium capture spots were subsequently labelled according to their overlap with the annotation polygons (**Supplementary Fig.4a**).

#### Validation with Xenium data

##### Correlation with reference data

Visium data was preprocessed by normalising (*pp.normalize_total*) and log1p-transforming (*pp.log1p*) the gene expression matrix using SCANPY. Once the ground truth data measured with Xenium were added, we discarded low-quality spots containing less than 1000 UMI counts (*pp.filter_cells*) and genes expressed in fewer than 10 spots (*pp.filter_genes*) using SCANPY. SVGs were first determined using Chrysalis with the following parameters: minimum Moran’s I=0.05, genes expressed in > 5% of the total number of capture spots. The detected SVGs were used to construct a low-dimensional representation using the dimensionality reduction step of Chrysalis. Chrysalis was subsequently used to define tissue compartments using archetypal analysis (number of PCs=20, number of compartments=8). To examine the relationship between the Xenium reference data and the compartments identified by Chrysalis, a correlation matrix was constructed by calculating Pearson correlation coefficients between these elements.

##### Marker gene contributions

SVG weights for each tissue compartment were calculated using Chrysalis to assess the importance of breast cancer and myoepithelial marker genes acquired from the original study^24^.

##### Assessing cell type composition of tissue compartments

To determine the cell type composition within each tissue compartment, we selected capture spots with domain scores above 0.8. We then aggregated the abundance data of the 20 mapped reference cell types and calculated their respective proportions per compartment.

#### Benchmarking

To benchmark methods based on their correlation with the Xenium reference data, we used the same pipeline described for the human lymph node dataset (see **Methods, Human lymph node analysis - Benchmarking** section).

#### Morphological feature extraction and integration

To incorporate morphological information, the spatial positions of capture spots from the Visium sample were first mapped to the 40X magnification post-Xenium H&E slide scan. Transformation matrices were used to translate the spatial positions between the two tissue sections (for details, see **Methods, Human breast cancer analysis - Co-registration of Visium and Xenium data** section). 3906 image tiles (299 × 299 px) were then extracted (**Supplementary Fig.5a**) by marking squares with centroids equal to those of the capture spots and side lengths equal to the capture spot diameters (55 µm). The image tiles were resized to 256 × 256 px, and the pixel values were standardised to be within the range of -1 to 1. The transformed images were used to train a deep autoencoder. The encoder was built using three 2D convolutional layers (kernel size: 3) and a dense layer, all coupled with GELU activation functions. The encoder output (i.e. bottleneck) is a 1-dimensional vector of 512. The decoder contains one dense layer and three 2D transposed convolution layers (kernel size: 3). The first three layers utilised GELU activation functions, while a Tanh activation function was employed for the last layer, resulting in an output size equivalent to that of the input image (pre-trained model is available at Zenodo). The architecture was built and trained using PyTorch^42^ and PyTorch-Lightning for 100 epochs with a batch size of 1 and learning rate decay using the Adam optimiser. After training, the latent vector for each encoded image tile was collected, resulting in a 2D matrix of 3906 × 512. Next, a PCA was performed on this matrix and the PCs were min-max scaled using scikit-learn’s *MinMaxScaler* function. Simultaneously, the same operation was applied to the PCs of the SVG expression matrix. The scaled morphology PCs (10) were then concatenated with the SVG expression PCs (20), and the resulting matrix of 30 PCs was subsequently processed with Chrysalis to find tissue compartments (number of PCs=30, number of compartments=8).

To demonstrate the benefits of the multimodal approach, we assessed the performance of Chrysalis relying on our pathologist’s annotations (see **Methods, Human breast cancer analysis - Mapping pathologist annotations to the Visium dataset**). We assigned every spot to the tissue compartment with the highest domain score (**Supplementary Fig.5b**). ARI was computed between those compartments and the annotation labels. F_1_ score was then calculated for every combination of annotations and tissue compartments computed by the gene expression-based and morphology-integrated approaches (**Supplementary Fig.5c**). Precision-recall curves for selected annotations were calculated using the respective tissue compartments with the highest F_1_ score (**Supplementary Fig.5d**).

### Mouse brain analysis

Anterior half of a mouse parasagittal brain section data was downloaded from the 10X Genomics online data repository. Manual annotations were acquired from another publication^21^. Capture spots containing less than 1000 UMI counts were omitted (*pp.filter_cells*), and genes expressed in 10 or fewer capture spots were also removed (*pp.filter_genes*) using SCANPY. This was followed by a normalisation (*pp.normalize_total*) and log1p-transformation (*pp.log1p*) step.

Chrysalis was used to determine SVGs with the following parameters: minimum Moran’s I=0.05, genes expressed in > 5% of the total number of capture spots. The detected SVGs were used to construct a low-dimensional representation using the dimensionality reduction step of Chrysalis. Chrysalis was used to define tissue compartments (number of PCs=20, number of compartments=[8, 12, 16, 20, 24, 28]) and to visualise them (**Supplementary Fig.7b**). Based on the results of 28 tissue compartments, the neocortex layers (Compartments 4, 15, 8, 1, and 11) were manually identified. The mean domain score of these compartments was calculated based on manual annotations obtained by consolidating the original annotations depicted in the upper panel of **Supplementary Fig.6b** for each cortical layer. The visual representation of the tissue compartments was examined and directly annotated based on the Allen Mouse Brain Atlas^28^. Additionally, we performed uniform manifold approximation and projection (UMAP) on the dataset using SCANPY. Chrysalis’s MIP method was used to shade the capture spots to verify the proximity of cortical tissue compartments in the UMAP space (**Supplementary Fig.7c**).

### Sample integration

Anterior and posterior parts of parasagittal mouse brain datasets were downloaded from the 10X Genomics online data repository. Capture spots containing less than 1000 UMI counts (*pp.filter_cells*) and genes expressed in 10 or fewer capture spots were removed (*pp.filter_genes*) using SCANPY. Data integration was performed with Scanorama^43^ after the initial normalisation (*pp.normalize_total*) and log1p-transformation (*pp.log1p*) of the count matrix for each sample. Following that, Chrysalis was used to calculate SVGs for both samples separately (minimum Moran’s I=0.20, genes expressed > 10% of the capture spots), and the SVG gene sets were concatenated. This resulted in 1325 SVGs with 675 genes overlapping between the two samples (**Supplementary Fig.8c**). The resulting SVG count matrix was used for dimensionality reduction and spatial domain detection by Chrysalis to identify tissue compartments (number of PCs=20, number of compartments=10).

### Slide-seqV2 analysis

Slide-seqV2 data of mouse hippocampus was downloaded using the link provided by the original study^3^. The count matrix and the corresponding spatial coordinates were read, and the appropriate AnnData data structure was manually constructed with SCANPY. Spatial locations beyond a circle with a radius of 2440 px from the mean XY spatial coordinate pair were removed to discard spots outside of the original ROI. The data was binned into square-shaped bins with a size of 24.4 px to create an evenly spaced array from the randomly dispersed capture beads. Bins containing less than two beads were omitted from further analysis. This was followed by removing capture spots containing less than 100 UMI counts (*pp.filter_cells*) and genes expressed in 10 or fewer bins (*pp.filter_genes*) using SCANPY. The count matrix was then normalised (*pp.normalize_total*) and log1p-transformed (*pp.log1p*). Chrysalis was used to find SVGs (minimum Moran’s I=0.05, genes expressed > 5% of the bins, number of neighbours for Moran’s I calculation=8). SVGs were used to construct a low-dimensional representation using the dimensionality reduction step of Chrysalis. Subsequently, Chrysalis was used to define tissue compartments using archetypal analysis (number of PCs=20, number of compartments=11).

### Stereo-seq analysis

Stereo-seq data of mouse embryo cross sections (E9.5, E12.5 - bin 50) were downloaded from MOSTA: Mouse Organogenesis Spatiotemporal Transcriptomic Atlas data repository. The gene expression matrix in these datasets is tiled up to 25×25 μm rectangle bins. HDF files were read with SCANPY, and bins labelled as cavities were removed using the annotations provided to the datasets. Normalisation (*pp.normalize_total*) and log1p-transformation (*pp.log1p*) were subsequently performed. Chrysalis was used to find SVGs (minimum Moran’s I=0.05, genes expressed > 5% of the bins, number of neighbours for Moran’s I calculation=8). SVGs were afterwards used to construct a low-dimensional representation using the dimensionality reduction step of Chrysalis. Then, Chrysalis was used to define tissue compartments using archetypal analysis (number of PCs=20, number of compartments=24).

## Data availability

10X Visium datasets used in this study, including the human lymph node, and the mouse brain anterior and posterior sections, are available on the 10X Genomics online data repository (https://www.10xgenomics.com/resources/datasets). 10X Xenium and Visium human breast cancer data and the corresponding high-resolution slide scans can be found on the 10X Genomics website (https://www.10xgenomics.com/products/xenium-in-situ/preview-dataset-human-breast). Cell type deconvolution data generated with cell2location for the human lymph node sample can be reproduced using the original pipeline (https://cell2location.readthedocs.io/en/latest/notebooks/cell2location_tutorial.html). Marker gene sets for cell types found in the lymph node are available on the CZ CELLxGENE Discover online resource (https://cellxgene.cziscience.com). Stereo-seq mouse embryo data can be accessed through the Mouse Organogenesis Spatiotemporal Transcriptomic Atlas (MOSTA) data repository (https://db.cngb.org/stomics/mosta). Slide-seqV2 mouse hippocampus data can be downloaded from the Broad Institute Single Cell Portal after registration (https://singlecell.broadinstitute.org/single_cell/study/SCP815/highly-sensitive-spatial-transcriptomics-at-near-cellular-resolution-with-slide-seqv2#study-summary).

AnnData objects for analysed datasets, all intermediary files, including cell type deconvolution results, marker gene sets for the human lymph node, the single cell labels for the Xenium data, reference data, and the expert annotations required to reproduce the analysis presented in this study are deposited at Zenodo (https://doi.org/10.5281/zenodo.8247780).

## Code availability

Chrysalis is available as a Python package at https://github.com/rockdeme/chrysalis. Documentation and tutorials are available at https://chrysalis.readthedocs.io. The code used for the data analysis in this article is available at https://github.com/rockdeme/chrysalis/tree/master/article.

## Acknowledgements

We thank M. Decollogny and S. Bagatella for providing guidance and assistance with the mouse brain tissue histology. This work was supported by the European Research Council (ERC-2019-AdG-883877), the Swiss Cancer League (KFS-5519-02-2022), and the Department of Defense (Award No. W81XWH-22-1-0557).

## Author Information

### Authors and Affiliations

**Institute of Animal Pathology, Vetsuisse Faculty, University of Bern, Bern, Switzerland** Demeter Túrós, Sven Rottenberg

**Roche Pharma Research and Early Development, Pharmaceutical Sciences, Roche Innovation Center Basel, F. Hoffmann-La Roche Ltd, Basel, Switzerland** Jelica Vasiljevic, Kerstin Hahn, Alberto Valdeolivas

**Bern Center for Precision Medicine (BCPM), University of Bern, Bern, Switzerland** Sven Rottenberg

### Contributions

D.T. conceived the idea of Chrysalis. A.V. and S.R. contributed to the study conceptualisation and supervised the research. D.T. designed the methodology, algorithms, validation and benchmarking analysis with input from S.R., J.V., and A.V. and implemented Chrysalis software. J.V. and D.T. developed morphological feature integration. K.H. performed histopathological validation and tissue annotations. D.T. wrote the original draft, J.V., K.H., A.V., S.R., and D.T. revised and edited the manuscript. S.R. worked on project administration and funding acquisition.

### Corresponding author

Correspondence to <u>Demeter Túrós</u>, <u>Alberto Valdeolivas</u>, or <u>Sven Rottenberg</u>.

