## Supplementary Materials for "Chrysalis: decoding tissue compartments in spatial transcriptomics with archetypal analysis"

#### Supplementary Figures

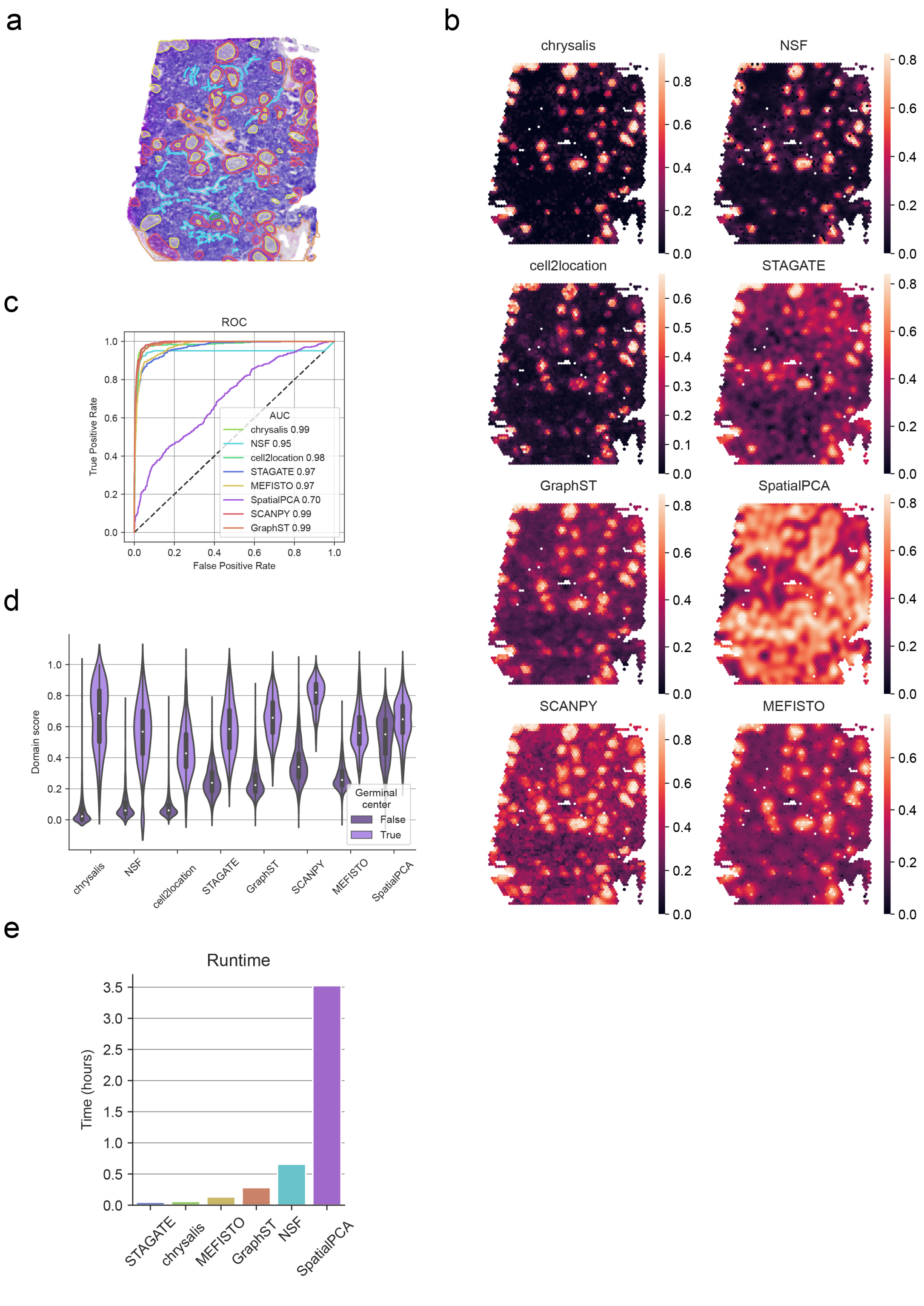

**Supplementary Fig.1 | Comparative analysis of spatial domain detection methods in identifying germinal centres**

**a,** Expert annotations for the different tissue types found in the human lymph node sample (yellow: germinal centres, red: mantle zone, light blue: large sinuses, orange: medulla, dark green: follicular tissue, not labelled: paracortex). **b,** Spatial domains corresponding to the germinal centres (rescaled to the range of 0-1) calculated using Chrysalis, NSF, cell2location (aggregated cell type abundances), STAGATE, GraphST, SpatialPCA, SCANPY, and MEFISTO. **c,** ROC-AUC curves of the spatial domains corresponding to the germinal centres. **d,** Violin plots displaying domain score distributions of the annotated germinal centres and the rest of the sample. **e,** Computational running times of the examined methods.

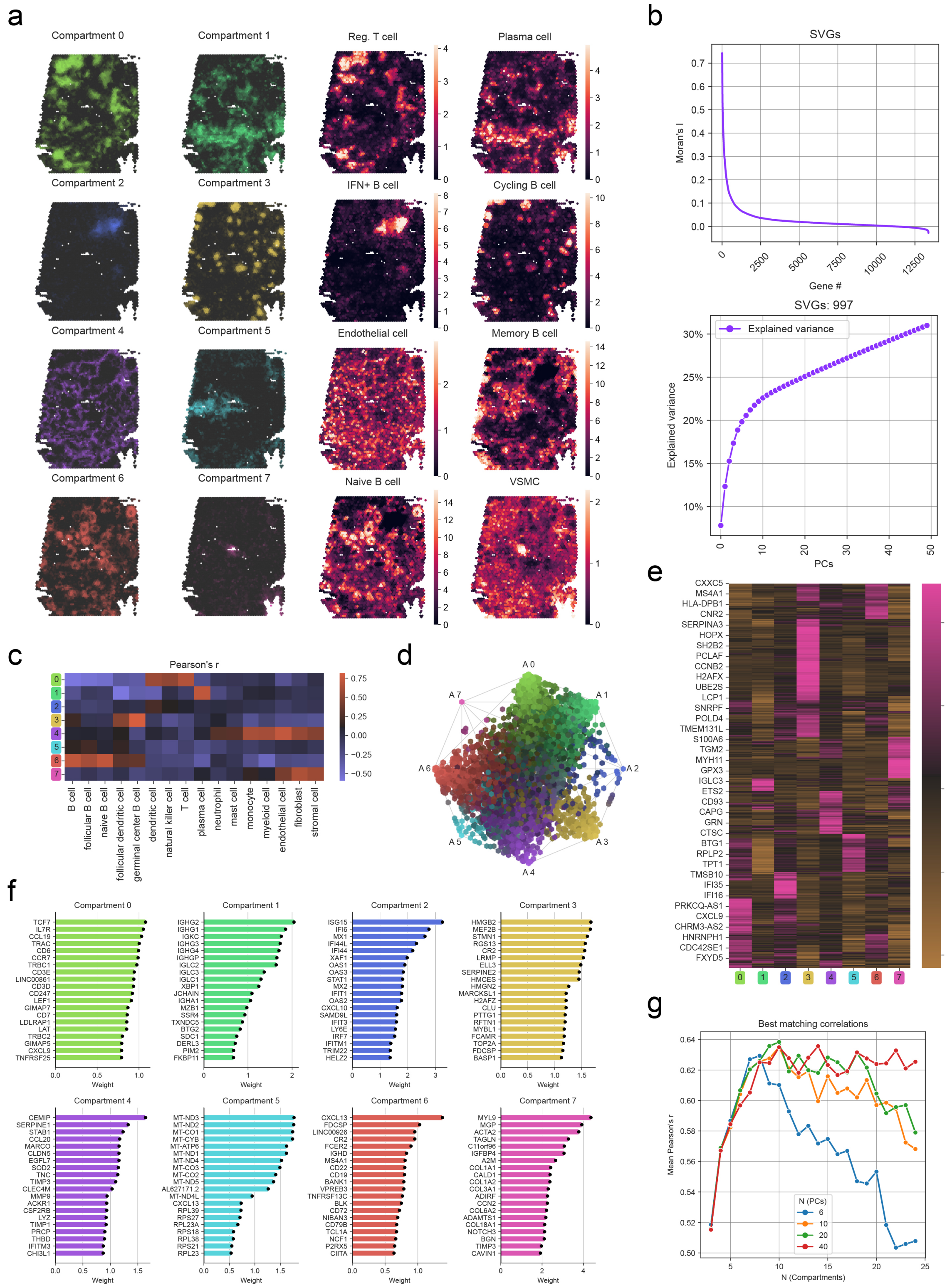


**Supplementary Fig.2 | Validation of Chrysalis in the human lymph node dataset**

**a,** Complete set of the compartments identified by Chrysalis (left panel) and highest correlating cell types from the deconvolution performed with cell2location (right panel). **b,** Rank-order plot of Moran's I for genes (upper panel) and explained variance of the low-dimensional embedding of the SVGs (lower panel). **c,** Correlation matrix showing Pearson's r values between the tissue compartments and the marker gene set expression of lymph node-specific cell types. **d,** Visualisation of the multi-dimensional simplex fitted to the low-dimensional embedding space showing the individual archetypes as the vertex points and the mixed tissue compartments represented by the individual colours of the remaining capture spots. **e,** Heatmap showing the contribution of SVGs to the identified tissue compartments. **f,** Top 20 contributing genes to all tissue compartments inferred by Chrysalis. **g,** Lineplot showing the changes in average correlation between the tissue compartments and the cell type deconvolution results with respect to tuning the number of PCs and archetypes.

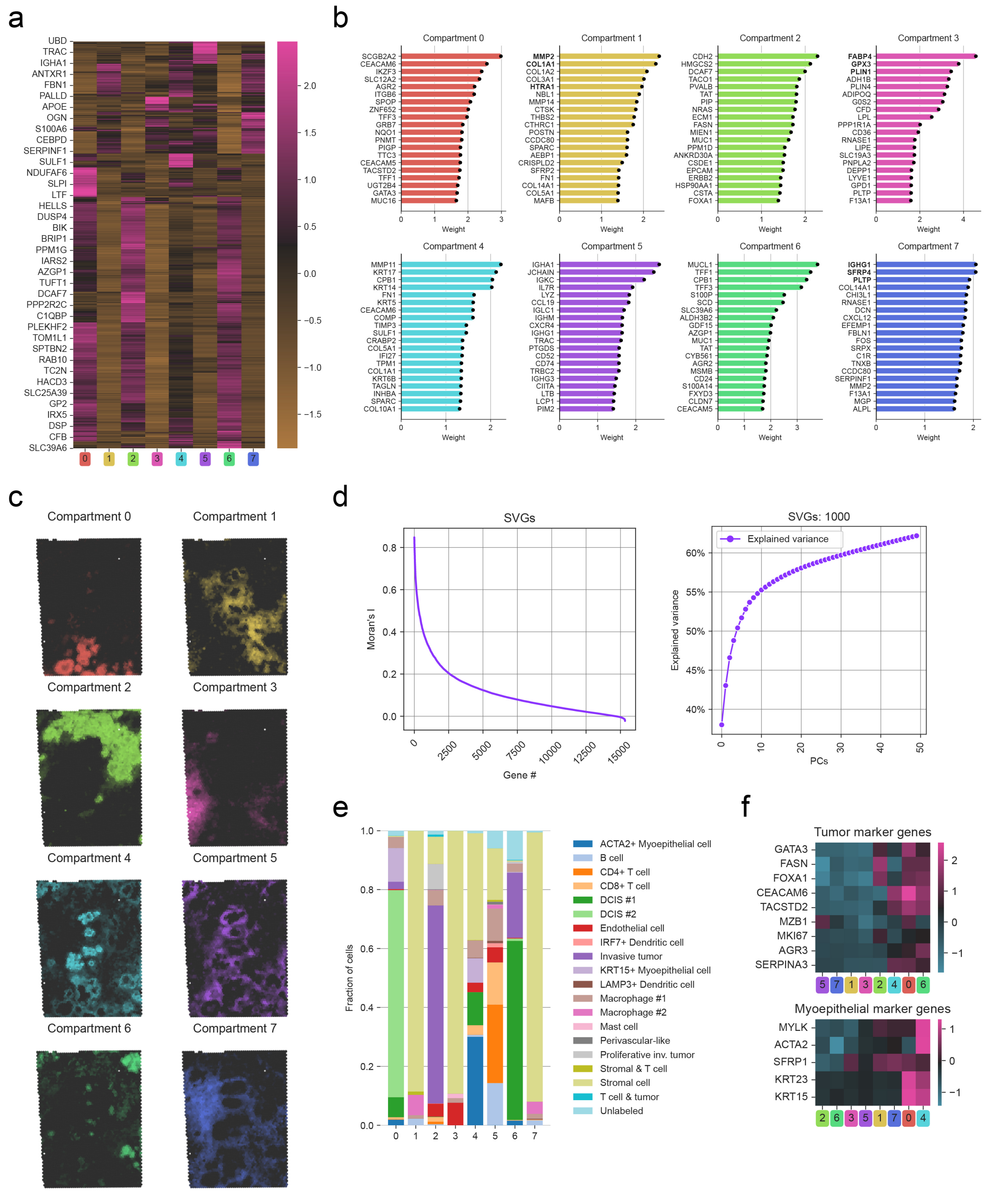


**Supplementary Fig.3 | Chrysalis identifies tissue compartments in the human breast cancer dataset**

**a,** Heatmap of SVG contributions to the identified tissue compartments. **b,** Top 20 contributing genes to tissue compartments. Bold characters were used to highlight the name of the genes referred to in the main text. **c,** Complete set of the individual tissue compartments. **d,** Rank-order plot of Moran's I for genes (left panel) and explained variance of the low-dimensional embedding of the SVGs (right panel). **e,** Cell type proportions of capture spots corresponding to pure tissue compartments (at least 0.8 domain score). **f,** Canonical tumour and myoepithelial marker gene weights for the compartments.

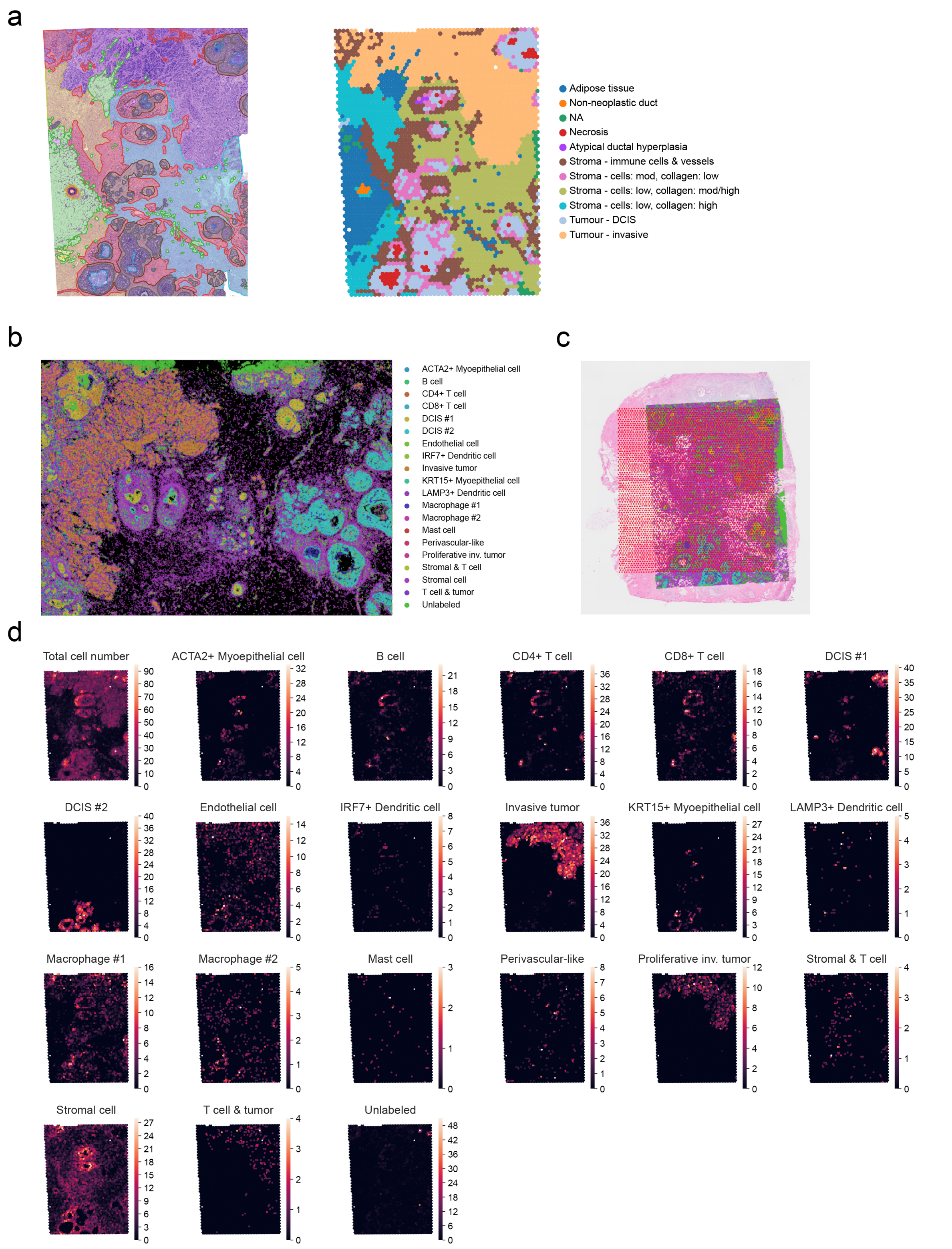


**Supplementary Fig.4 | Constructing reference for the human breast cancer dataset**

**a,** Expert annotations (left panel) for the different tissue types found in the post-Xenium H&E of the human breast cancer sample (green: *adipose tissue*, orange: *non-neoplastic duct*, blue: *necrosis*, pink: *atypical ductal hyperplasia*, red: *stroma - immune cells & vessels*, brown: *stroma - cells: mod, collagen: low*, light blue: *stroma - cells: low, collagen: mod/high*, yellow: *stroma - cells: low, collagen: high*, grey: *tumour - DCIS*, purple: *tumour - invasive*) and the Visium capture spots corresponding to these regions (right panel) after the alignment. **b,** Distinct cell types in the Xenium data (single cell centroids depicted with various colours). **c,** Alignment of Xenium (single cell centroids depicted with various colours) and Visium data (red hexagonal grid) overlaid on top of the pre-Visium H&E image showing the area of overlap between the two modalities. **d,** Aggregated ground truth cell type abundance data for each capture spot.

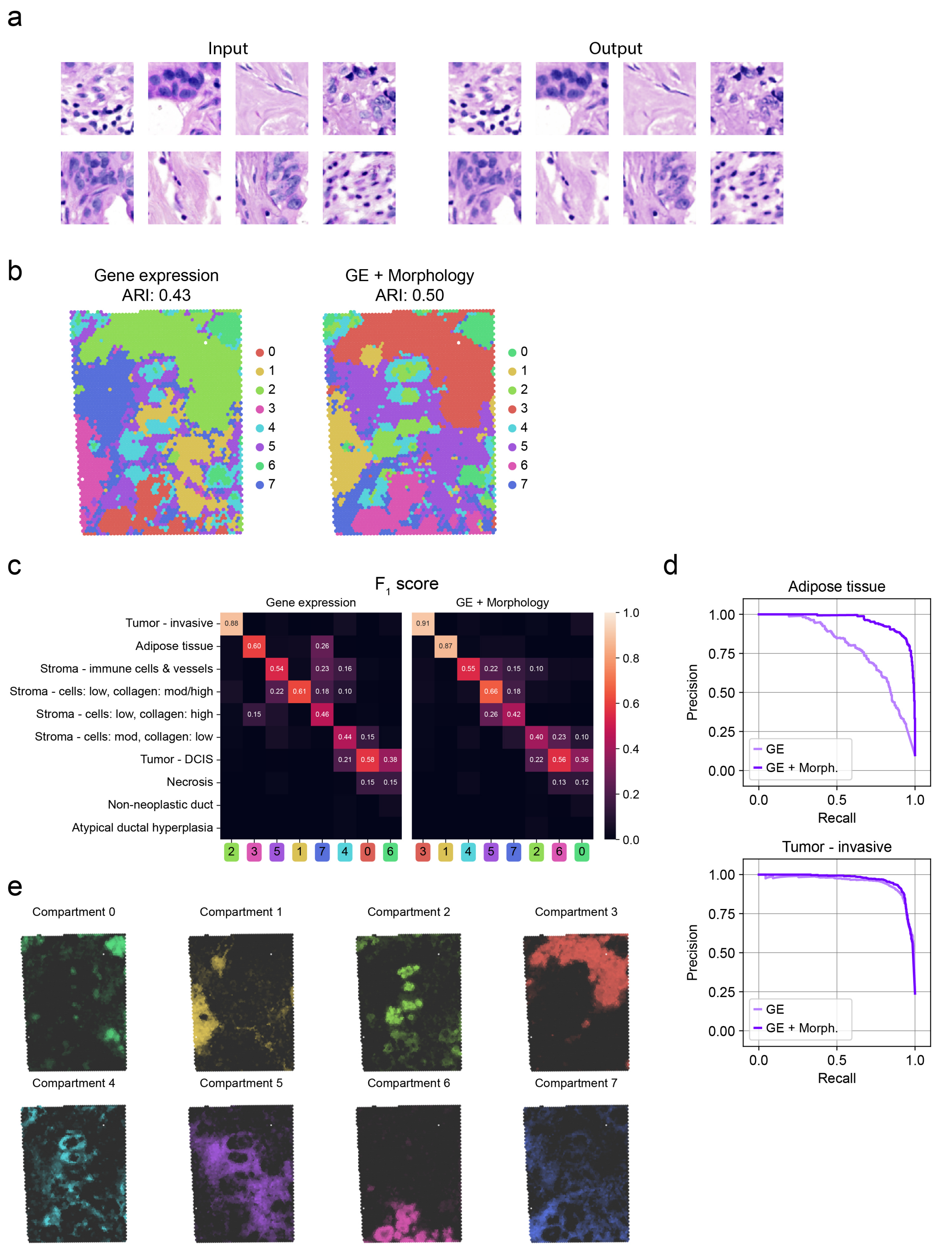


**Supplementary Fig.5 | Morphological feature integration with Chrysalis**

**a,** Autoencoder input and output image tiles from the post-Xenium H&E of the human breast cancer sample. Each image tile shares its centroid with the corresponding capture spot and has the same side length as the diameter of capture spots (55 μm). Output image tiles show that the autoencoder can successfully reconstruct the input images using only the information encoded in the bottleneck. **b,** Each capture spot is labelled with the tissue compartment associated with the highest domain score based on the gene expression-based (left panel) and morphology-enhanced (right panel) results. **c,** Heatmaps showing $F_1$ score values for each tissue compartment-annotation pair using the gene expression-based and morphology-enhanced low-dimensional embedding (values < 0.10 are not displayed). **d,** Precision-recall curves for two selected annotations. Curves of tissue compartments with the highest $F_1$ scores are shown for each annotation (GE: gene expression-based tissue compartments, GE + Morph.: morphology-enhanced tissue compartments). **e,** Individual tissue compartments inferred by Chrysalis using the morphology integrated low-dimensional embedding.

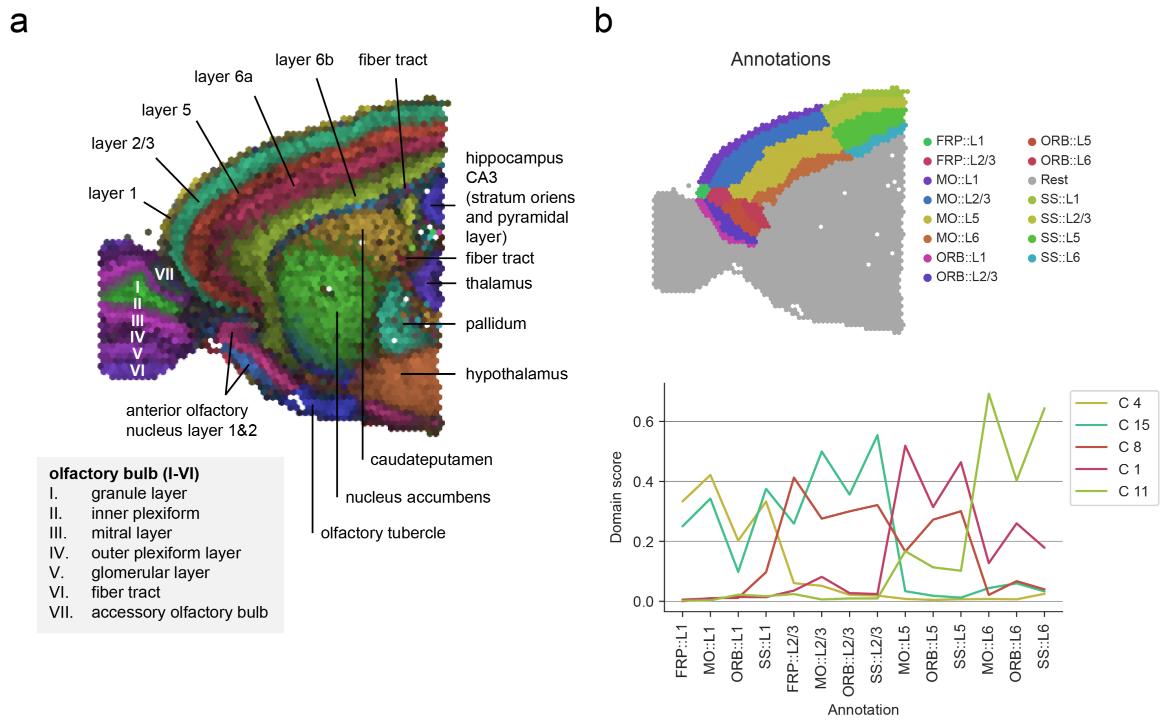


**Supplementary Fig.6 | Validation of Chrysalis using expert annotations on the mouse brain anterior sample**

**a,** Direct annotation of Chrysalis's tissue compartments based on the Allen Mouse Brain Atlas. **b,** Expert annotations (top panel) and the corresponding mean domain scores of the selected compartments (bottom panel).

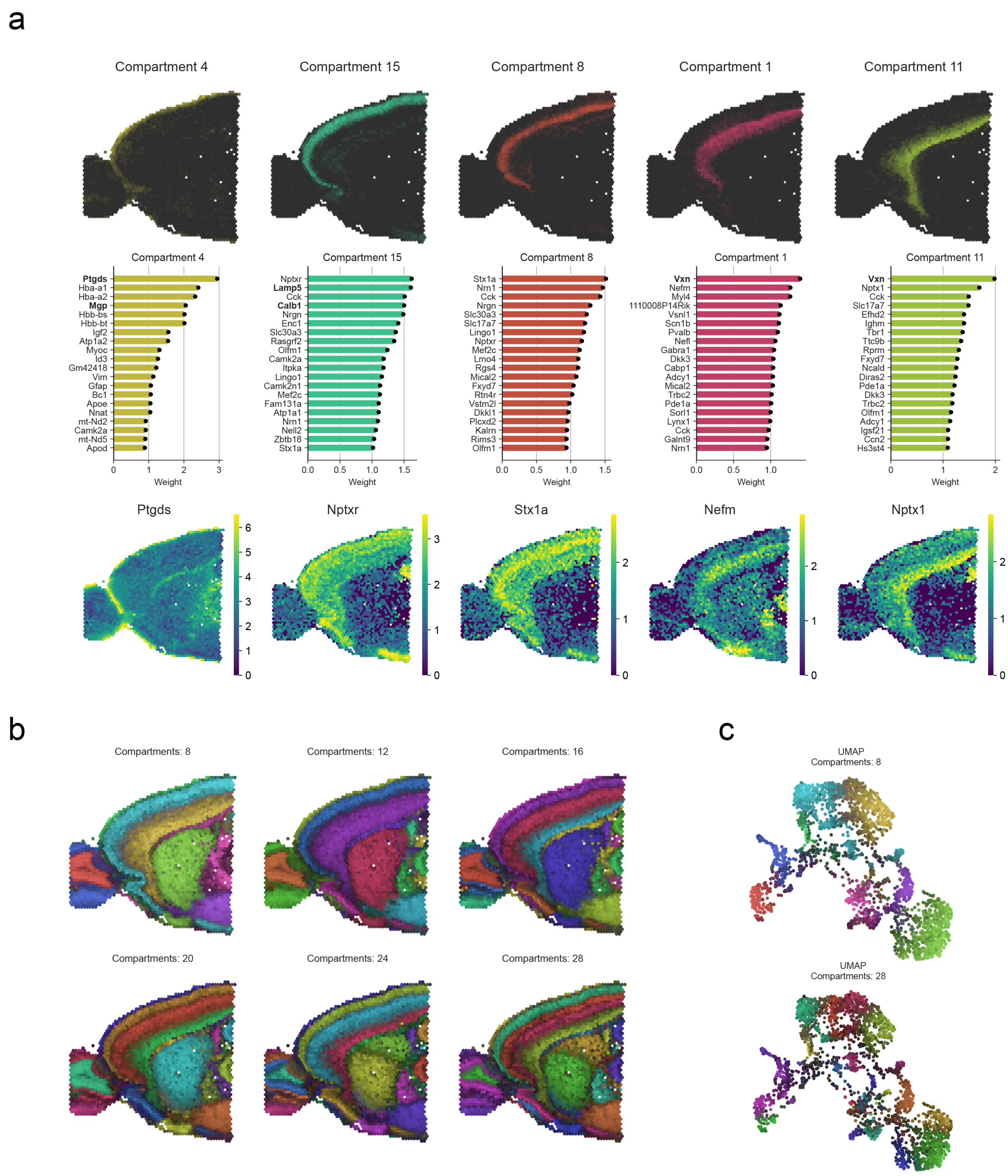


**Supplementary Fig.7 | Analysis of anatomical regions in the mouse brain anterior sample inferred by Chrysalis**

**a,** Five tissue compartments representing layers 1, 2/3, 4, 5 and 6 (from left to right). Each compartment is accompanied by the top 20 genes weighted according to their contribution to that specific compartment. The lower panel illustrates the expression profile of genes with the highest weights. Bold characters were used to highlight the name of the genes referred to in the main text. **b,** Chrysalis MIP of the mouse brain with 8, 12, 16, 20, 24, and 28 distinct compartments. **d,** UMAP of the mouse brain data, capture spots are coloured according to the corresponding MIP with 8 and 28 compartments.

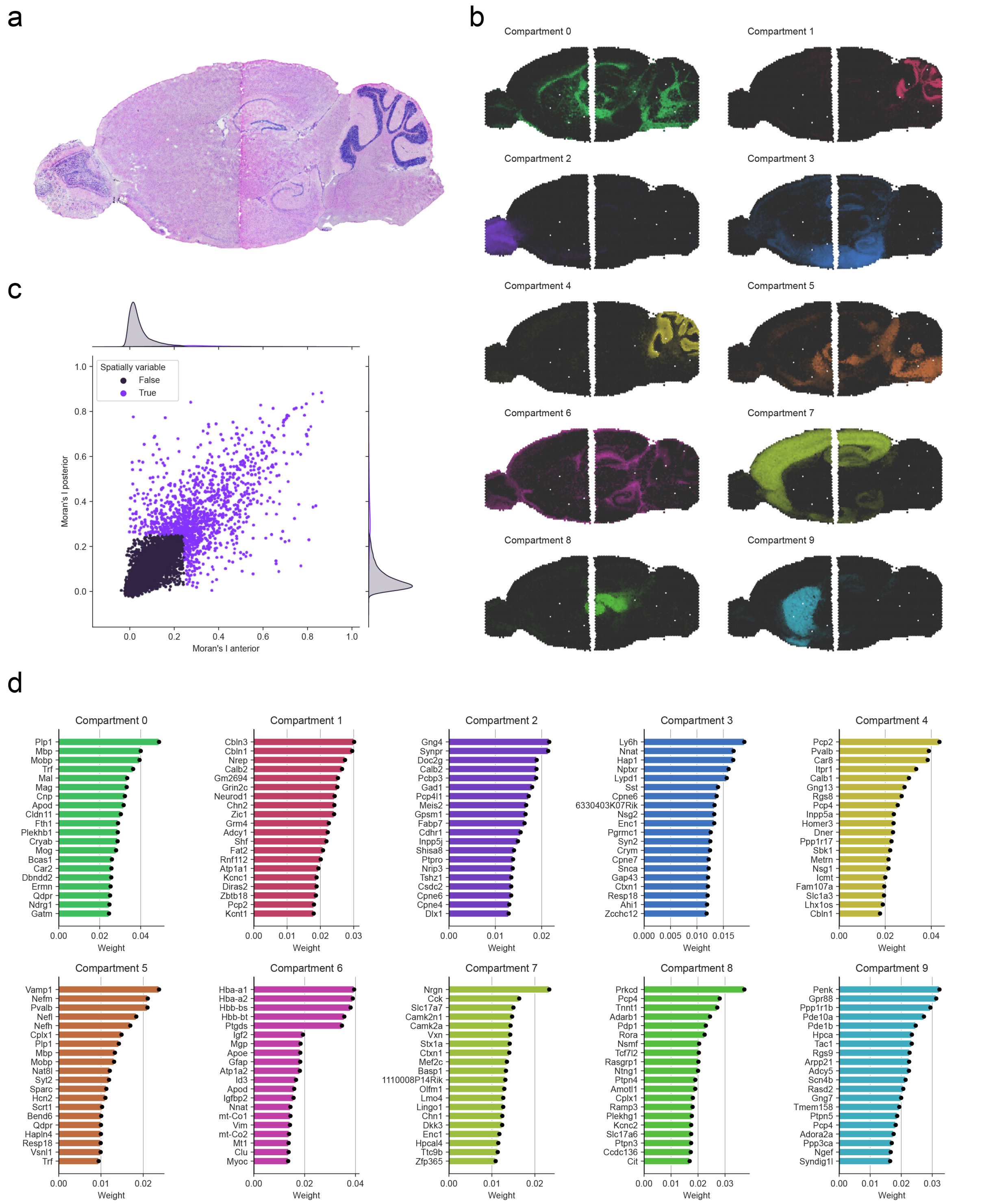


**Supplementary Fig.8 | Multi-sample dataset**

**a,** Mouse brain anterior and posterior H&E images. **b,** Complete set of the individual tissue compartments inferred by Chrysalis. **c,** Moran's I values for genes across the anterior and posterior sections (identified SVGs labelled with light purple). **d,** Top 20 contributing genes to tissue compartments.

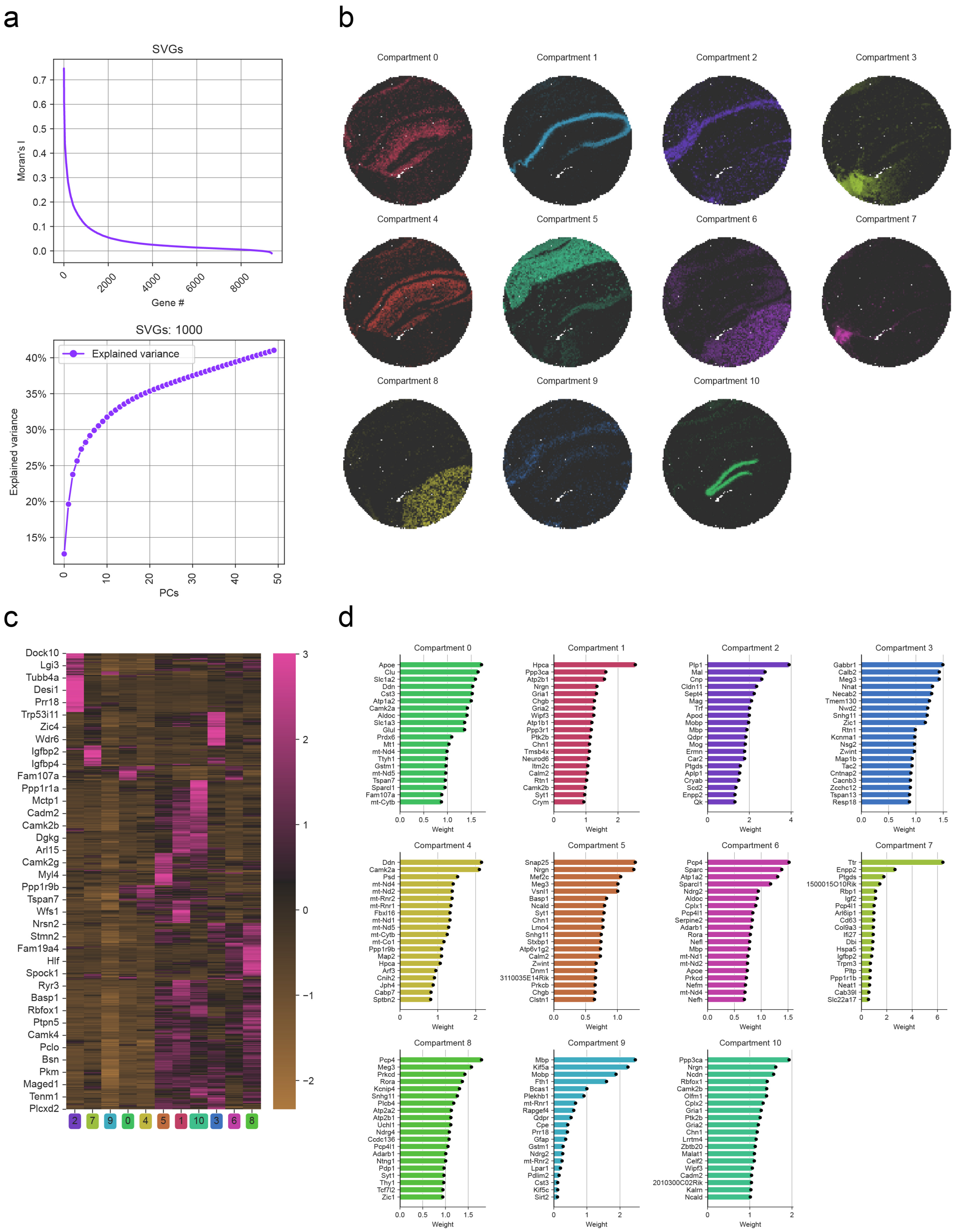


**Supplementary Fig.9 | Tissue compartments in the Slide-seqV2 mouse hippocampus dataset**

**a,** Rank-order plot of Moran's I (upper panel) and explained variance of the low-dimensional embedding of the SVGs (lower panel). **b,** Complete set of the individual Chrysalis compartments. **c,** Heatmap of SVG contributions of the identified tissue compartments. **d,** Top 20 contributing genes to tissue compartments predicted by Chrysalis.

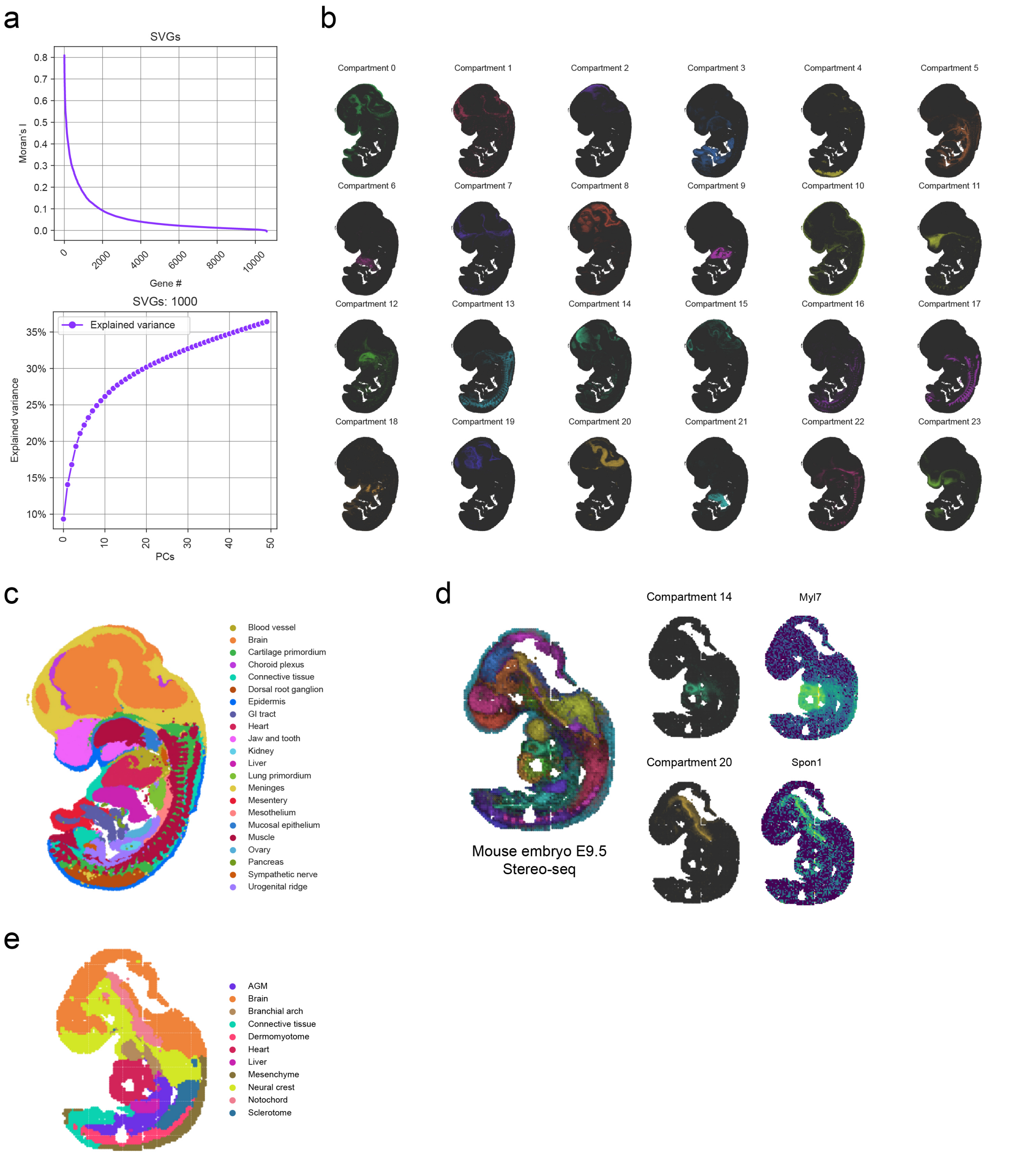


**Supplementary Fig.10 | Tissue compartments in the Stereo-seq mouse embryo datasets**

**a,** Rank-order plot of Moran's I (upper panel) and explained variance of the low-dimensional embedding of the SVGs (lower panel) for the E12.5 sample. **b,** Top 20 contributing genes to tissue compartments predicted by Chrysalis for the E12.5 sample. **c,** Tissue type annotations for the E12.5 sample from the original publication. **d,** Chrysalis MIP in the E9.5 sample (left panel) and the compartments corresponding to the heart (upper right panel) and the notochord (lower right panel) with their respective top-weighted genes. **c,** Tissue type annotations for the E9.5 sample from the original publication.

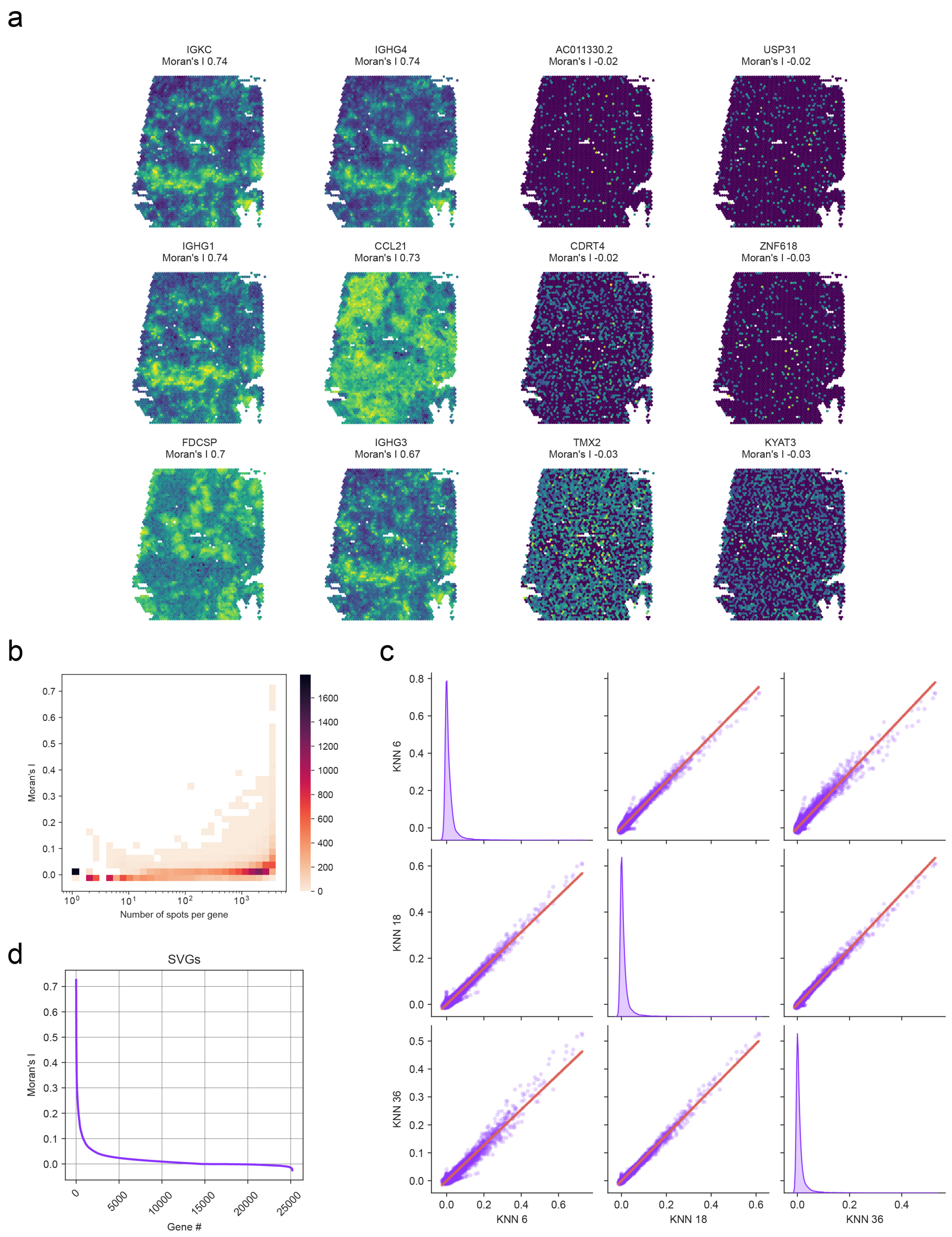


**Supplementary Fig.11 | Detecting spatial autocorrelation with Moran's I**

**a,** Spatial gene expression profile of genes with the 6 highest and lowest Moran's I value in the human lymph node dataset. **b,** 2D heatmap of the distribution of Moran's I values in function of the number of capture spots the respective genes were detected in. **c,** Pairplot showing the impact of using KNN 6, 18 and 36 on Moran's I statistic. **d,** Rank-order plot of

Moran’s I for all genes found in the human lymph node sample (without omitting the sparsely expressed genes).

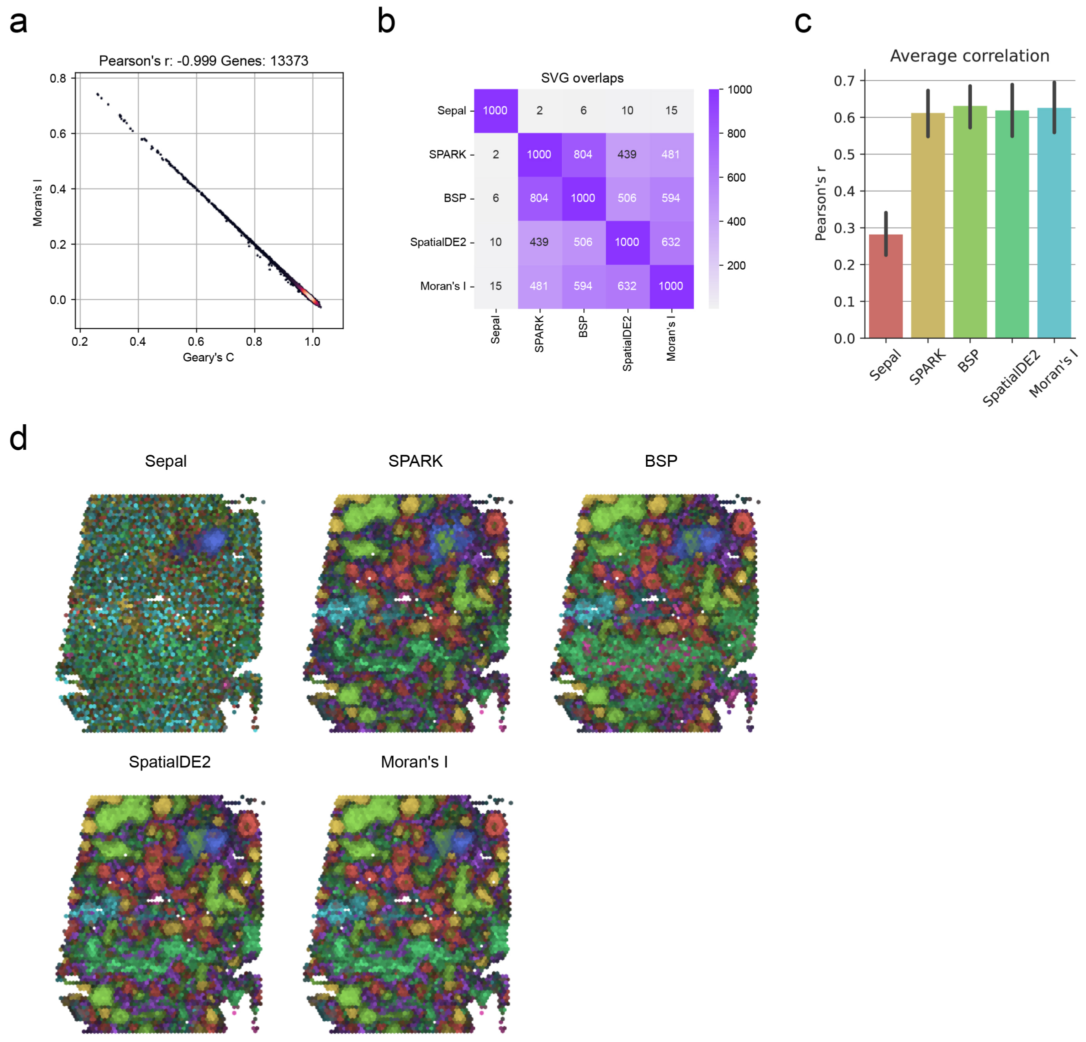


**Supplementary Fig.12 | Comparison of SVG detection methods in the human lymph node dataset**

**a,** Comparison of Moran's I and Geary's C statistics. **b,** Overlap between the top 1000 SVGs across the examined methods. **c,** Correlation of cell type deconvolution results with the Chrysalis compartments using the SVGs determined by the examined methods (bar heights denote the mean correlation coefficient, and error bars show standard deviation). **d,** Chrysalis's MIP of tissue compartments identified using different SVG detection algorithms.

### Supplementary Tables

#### SVG detection methods

| Name | Tutorial | GitHub |
|---|---|---|
| SpatialDE2 | https://github.com/PMBio/spatialde2-paper | https://github.com/PMBio/SpatialDE |
| SEPAL | https://squidpy.readthedocs.io/en/stable/notebooks/examples/graph/compute_sepal.html | https://github.com/almaan/sepal |
| BSP | https://github.com/juexinwang/BSP/blob/main/README.md | https://github.com/juexinwang/BSP |
| SPARK-X | https://xzhoulab.github.io/SPARK/02_SPARK_Example | https://github.com/xzhoulab/SPARK |

#### Benchmarking

| Name | Tutorial | GitHub |
|---|---|---|
| NSF | https://github.com/willtownes/nsf-paper/tree/main/scrna/visium_brain_sagittal | https://github.com/willtownes/nsf-paper |
| MEFISTO | https://github.com/bioFAM/MEFISTO_tutorials/blob/master/MEFISTO_ST.ipynb | https://github.com/bioFAM/mofapy2 |
| STAGATE | https://stagate.readthedocs.io/en/latest/T1_DLPFC.html | https://github.com/zhanglabtools/STAGATE |
| SpatialPCA | https://lulushang.org/SpatialPCA_Tutorial/DLPFC.html | https://github.com/shangll123/SpatialPCA |
| GraphST | https://deepst-tutorials.readthedocs.io/en/latest/Tutorial%201_10X%20Visium.html | https://github.com/JinmiaoChenLab/GraphST |
| SCANPY | https://scanpy-tutorials.readthedocs.io/en/latest/spatial/basic-analysis.html | https://github.com/scverse/scanpy |
| cell2location | https://cell2location.readthedocs.io/en/latest/notebooks/cell2location_tutorial.html | https://github.com/BayraktarLab/cell2location |

**Supplementary Table S1 | List of computational methods used for SVG detection and spatial domain inference**
